# Effects of EEG Preprocessing on Channel-Wise Attention and Effective Connectivity Alignment in Visual EEG Decoding

**DOI:** 10.64898/2026.07.02.736026

**Authors:** Virginia Valensiene Elichatiti, Basari, Muhammad Arif, Mohammad Ikhsan

**Affiliations:** Department of Electrical Engineering, Universitas Indonesia, Depok, Jawa Barat 16424, Indonesia; Research Center for Biomedical Engineering, Universitas Indonesia, Depok, Jawa Barat 16424, Indonesia; Department of Molecular and Clinical Medicine, University of Gothenburg, The Wallenberg Laboratory, Sahlgrenska University Hospital, Gothenburg, Sweden

**Keywords:** Electroencephalogram, preprocessing, attention, connectivity, brain-computer interface (BCI)

## Abstract

Transformer-based deep learning models have shown great potential for decoding visual EEG signals. However, their internal attention mechanisms are often evaluated primarily on optimization objectives, leaving their alignment with biological brain connectivity an open question. This study empirically evaluates how variations in EEG preprocessing strategies affect these attention representations using the Adaptive Thinking Mapper (ATM) model as a framework. We compared a baseline pipeline (MVNN only) against a comprehensive cleaning pipeline integrating ICA and notch filtering. The models were evaluated through cross-generalization, noise robustness, and spectral-temporal ablation analyses. Furthermore, we investigated the structural correspondence between the model’s data-driven attention weights and neurophysiological reference networks (GPDC, PDC, and DTF) using Node Strength Correlation and Representational Similarity Analysis (RSA). The results show that the comprehensive preprocessing successfully suppresses non-neural artifacts, such as frontal noise and electrical interference, while maintaining comparable decoding accuracy and baseline robustness. Alignment analyses revealed that the broad spatial organization of the learned attention patterns remains highly stable across pipelines, capturing key directed connectivity dynamics with subtle, metric-dependent variations in global representational geometry. This work provides an empirical exploration into bridging data-driven attention weights with neurophysiological consistency, offering insights toward more transparent brain-computer interfaces.

## I. Introduction

THE rapid evolution of artificial intelligence (AI) technology has paved the way for the fields of computational neuroscience and brain-computer interface (BCI). In recent years, studies have gradually shifted from conventional methods to Transformer-based deep learning architecture and Generative AI. These methods aim to map abstract electroencephalogram (EEG) signals into a rich semantic feature space for visual image reconstruction [1], [2].

The research conducted by Li et al. introduced the Adaptive Thinking Mapper (ATM) model to encode EEG signals into embeddings [2]. The ATM architecture includes the channel-wise attention layer that functions to learn the relevance between EEG channels. Attention works by assigning weights to the connections between channels to optimize the objective function of the model [3]. Although this approach can produce accurate image reconstructions using a diffusion model from the embeddings, the model remains difficult to interpret biologically [4]. Attention in Transformer is data-driven and optimized based on loss functions. It does not explicitly incorporate causality or brain frequency dynamics [5].

However, biological interpretability is related not only to the suitability of brain connectivity patterns but also to how model representations are influenced by preprocessing strategies and signal spectral characteristics. Processes such as Independent Component Analysis (ICA), multivariate normalization (MVNN), and variations in certain frequency bands have the potential to alter the structure of representations learned by the model. In addition, temporal components such as event-related potential (ERP) responses in visual paradigms can also influence attention patterns. Therefore, a comprehensive analysis is needed to understand how preprocessing, frequency dynamics, and temporal characteristics shape attention representation in Transformer models.

In the context of visual processing, dynamic interactions between brain areas play an important role in integration and information transfer. Pathways like occipital-to-temporal and occipital-to-parietal show directional communication patterns [6]. Therefore, directed connectivity is important for understanding how information is processed and distributed in the brain. Effective connectivity metrics such as partial directed coherence (PDC), generalized PDC (GPDC), and directed transfer function (DTF) are based on multivariate autoregressive model (MVAR) to estimate directed interactions in the frequency domain [7]. This approach provides reference effective connectivity measures to evaluate whether the attention patterns learned by the model are consistent with the directed connectivity studied in the literature.

Drawing from the established impact of EEG artifact removal and the known neurophysiological markers of visual perception, this study evaluates three primary hypotheses. First, we hypothesize that while the proposed preprocessing pipeline will effectively suppress non-neural interference, specifically frontal ocular artifacts and line-noise, it will maintain comparable decoding accuracy and model robustness against signal noise compared to baseline methods. Second, we anticipate that the decoding capability of the model will be mainly based on low-frequency EEG components and early post-stimulus intervals from 0 to 400 ms, reflecting the primary role of initial sensory processing and attentional involvement in visual tasks. Finally, we hypothesize that a more refined preprocessing approach may influence the biological plausibility of the learned attention patterns by altering their structural correspondence with biologically motivated reference connectivity measures.

Building on this framework, the primary goal of this study is to systematically evaluate channel-wise attention representations in Transformer models for visual EEG decoding, specifically examining how preprocessing strategies affect their biological interpretability. To achieve this, our analysis is conducted through three main approaches:

1. comparison of preprocessing strategies,
2. comprehensive model evaluation through cross-generalization, spectral-temporal ablation, and noise robustness studies, and
3. evaluation of the alignment between the attention matrix and the effective connectivity matrix using node strength correlation and representational similarity analysis (RSA).

## II. Related Works

### A. EEG-Based Visual Decoding and Image Reconstruction

Research on image reconstruction from EEG signals has rapidly advanced through the use of deep learning approaches. Several studies have been conducted previously and have contributed to the development of methods for reconstructing images based on brain signals. Brain2Image is a framework that converts EEG signals into images using a deep learning model that shows potential for translating neuronal activity into a visual image [8]. This line of work was further advanced by ThoughtViz, which employed a Generative Adversarial Network (GAN) to generate high-quality images from EEG data [9]. Subsequently, a new method called Contrastive Learning-Image Pretraining (CLIP) emerged, opening new opportunities for further research by utilizing text-to-image capabilities in the visual reconstruction process from EEG signals [10]–[13]. Then, recent research attempted to develop a visual reconstruction method from EEG signals to match the quality of visual reconstruction from fMRI [2]. These approaches successfully improved the quality of reconstruction and semantic correspondence between EEG and target images. However, most studies focus on reconstruction performance without evaluating whether the model’s internal representation corresponded to neurophysiological mechanisms.

### B. Attention Mechanism

The attention mechanism is used as an adaptive weighting system that allows the model to prioritize the most relevant information from complex EEG data. Specifically, channel-wise attention can help reduce noise by assigning greater weight to channels that contribute more strongly to the decoding objective [2]. While attention can be associated with interpretability, attention weights are optimized by the training objective and do not explicitly represent neural causality [5]. In the context of EEG, attention measures the importance of a feature based on the mathematical objective of the loss function. Therefore, there is a risk that the model will assign high weights to artifacts or statistical patterns that have no functional meaning in the brain [14]. For this reason, thorough empirical evaluation is needed to determine how attention weights are consistent with neural interactions measured by conventional neurophysiological methods.

### C. Directed Connectivity in EEG

In neuroscience, interactions between brain regions can be measured using various modalities, including structural connectivity (SC), functional connectivity (FC), and effective connectivity (EC). SC is a physical representation of nerve pathways or brain fibers that connect various regions anatomically. Meanwhile, EC explains causal relationships that show how one area of the brain directly influences another area in a targeted manner. On the other hand, FC measures how synchronized the activity between two brain areas is based on similarities in statistical patterns without considering the direction of the relationship [7], [15].

In this research, EC is used because it can represent the directed influence between brain regions and the dynamic flow of neural information. In EEG analysis, EC is generally estimated with MVAR to model the causal relationship between channels in frequency domain. Within this framework, several widely used metrics are PDC, GPDC, and DTF, which are designed to measure directed interactions between EEG signals in a frequency-specific manner. This approach makes EC relevant for evaluating whether the attention weights in the Transformer model reflect the pattern of directed neural information flow [7], [15].

### D. Preprocessing Effects in EEG Data

EEG signals are very weak and often remain noisy even after physical filtering. Therefore, preprocessing is an important step to ensure that the signals are processed correctly and to avoid errors in interpretation [16].The main challenge in recording EEG signals is blinking, especially for experiments related to vision. Various types of artifacts, including blinking, can be overcome using ICA decomposition [17]. Numerous studies have shown that ICA improves signal quality and enhances downstream decoding performance [18]. However, aggressive component rejection may also alter the intrinsic spectral structure of EEG signals, particularly in low-frequency bands that are relevant for event-related potentials and large-scale cortical interactions [19]. Since effective connectivity metrics and attention mechanisms both rely on inter-channel relationships, changes introduced during preprocessing may influence the learned representations and their biological interpretation. Therefore, understanding how preprocessing strategies such as ICA affect attention representations is an important yet underexplored aspect in EEG-based deep learning models.

## III. Methods

### A. Dataset

The study was conducted on the THINGS-EEG2 dataset, which involved 10 healthy subjects and was acquired using the Rapid Serial Visual Presentation (RSVP) paradigm. In this research, data from all 10 subjects were utilized. This dataset consists of 16,540 training images and 200 testing images. Neural data were captured using a 63-channel EEG system that re-referenced to Fz electrode and with a sampling rate of 1000 Hz [20].

### B. Data Preprocessing

The preprocessing pipeline began with cleaning and artifact removal from the raw data (63-channel, 1000 Hz). There are 63 channels instead of 64, as the data have been re-referenced to the Fz electrode.Our initial step was to apply a 50 Hz notch filter to suppress power-line interference and its associated harmonics, which could otherwise obscure subtle neural oscillations. Next, we applied ICA decomposition with 63 components to isolate and remove physiological artifact components, such as blinking and muscle activity, without damaging the relevant brain signals. Artifact-related ICA components were identified based on their characteristic temporal profiles and scalp topographies, with particular attention to ocular and muscle-related patterns. After the signal was cleaned, we carried out the data segmentation stage, in which the signal was cut in a time range of -0.2 seconds to 1.0 seconds relative to the stimulus onset, followed by baseline correction and downsampling to 250 Hz for computational efficiency. Specifically, for deep learning model input preparation, the MVNN technique is applied through covariance matrix calculation and whitening processes to normalize the noise structure between channels before the data is fed into the Transformer architecture. We also compare our pipeline with the literature method [2] that only applied the MVNN technique without involving the notch filtration or ICA decomposition processes as the baseline in this study. The preprocessing flow can be seen in Fig. 1A.

**Fig. 1.**
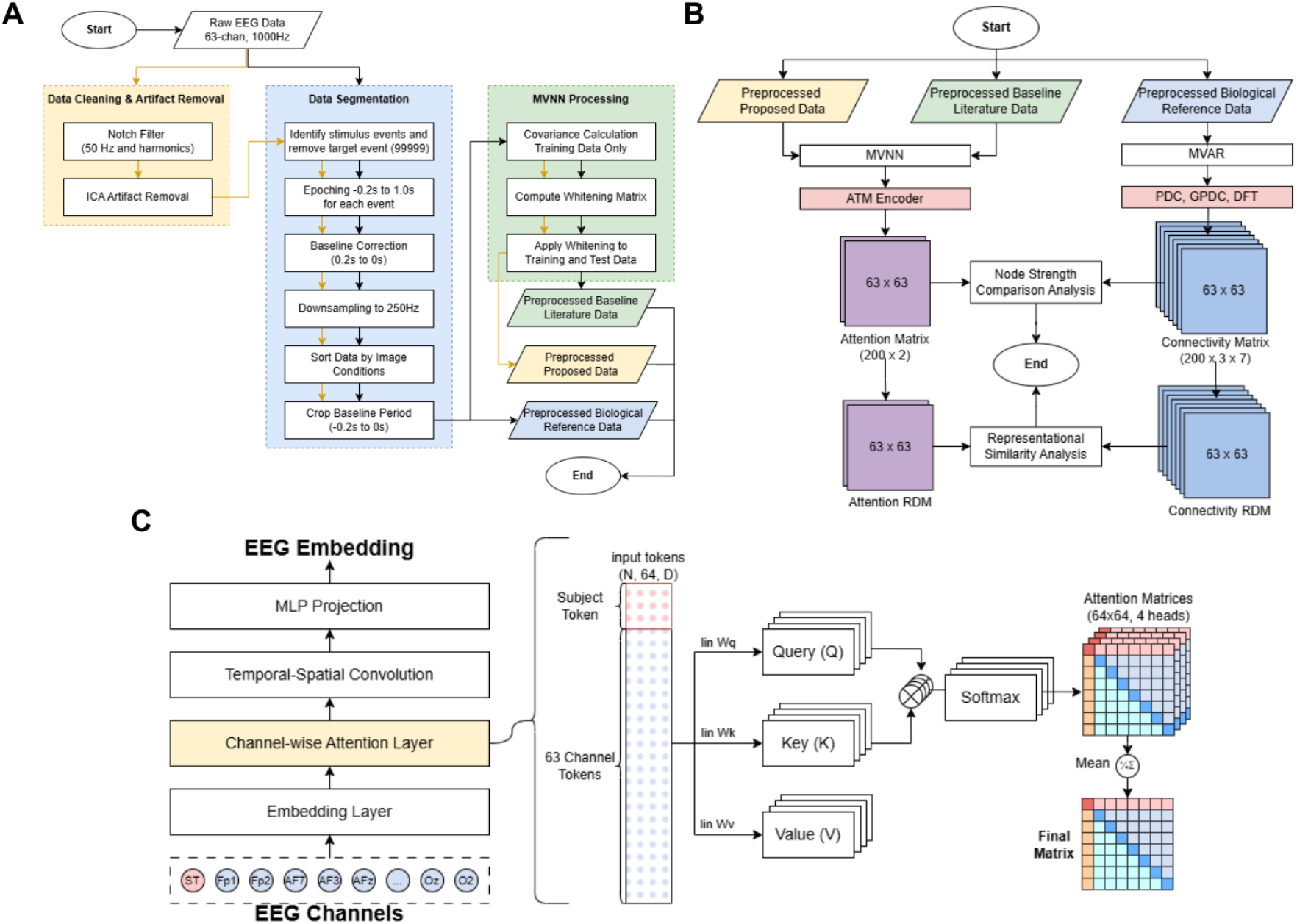
Overall methodology and architecture framework. (A) The EEG preprocessing workflow, comparing the baseline pipeline with the proposed pipeline (incorporating notch filtering, ICA, and MVNN). (B) The analytical framework for evaluating the alignment between the model’s attention matrix and the reference effective connectivity using Node Strength Comparison and Representational Similarity Analysis (RSA). (C) The Adaptive Thinking Mapper (ATM) encoder architecture, highlighting the extraction of the channel-wise attention matrix from the multi-head self-attention layer [2].

### C. Model Architecture

The model used in this study is based on a Transformer architecture that has been specifically modified for EEG signal processing. The architecture is designed to align the EEG representations with the CLIP latent space, essentially training the model to mirror the semantic “logic” of CLIP for use in neural decoding. This mapping appears necessary to translate complex temporal dynamics into a structured visual-semantic framework suitable for image reconstruction. However, in this study, we focus on the Channel-wise Attention Layer from the encoder (Fig. 1C) [2].

The input consists of Subject Token that capture inter-subject variability, which appears to help the model remain adaptable to the unique signal characteristics of individual participants. Alongside this, 63 Channel Tokens are incorporated to directly represent the primary signal data from the corresponding EEG electrodes. The data is then processed through the multi-head self-attention mechanism that allows the model to learn relationships between channels. The attention score is formulated as:

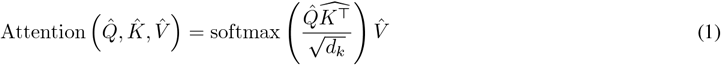

In the process, the input is projected through a linear layer and divided into four parallel heads to study various feature spaces (subspaces) of the representation simultaneously before being recombined for final analysis [2].

### D. Model Performance Evaluation

The model was optimized using AdamW with a learning rate of 3e-4 and a batch size of 64. To ensure complete reproducibility of the training dynamics and data shuffling, the random seed was fixed at 24 across all experiments. The reported mean and standard deviation (SD) for classification accuracy were calculated across the 10 subjects. Each model was trained for 40 epochs under identical optimization settings to ensure consistency across experimental conditions. Evaluation was performed using the final epoch model rather than selecting the best-performing checkpoint to avoid potential selection bias and to enable a fair comparison of learned representations and generalization behavior. Model performance was assessed using classification accuracy on the testing set, reported in terms of Top-*k* accuracy. Top-1 accuracy reflects the standard metric in which only the highest-probability prediction is considered correct, while Top-5 accuracy measures whether the ground-truth label appears among the five most confident predictions, providing a more flexible evaluation of ranking quality [2].

In addition to Top-*k* metrics, performance was also analyzed under different candidate-set settings, namely V2, V4, and V10. In the V2 configuration, the model selected the correct image from two candidates. In V4, the selection was made from four candidates. In V10, the model performed classification among ten candidates, reflecting the increasing levels of task difficulty. These configurations provide insight into the robustness of the model under varying levels of ambiguity [2]. However, the primary analysis focuses on Top-1 and Top-5 accuracy as the main indicators of classification performance.

To further evaluate cross-condition generalization, a cross-test experiment was conducted [21], [22]. Specifically, the model trained on baseline-preprocessed data was evaluated using test data processed with the proposed pipeline, and conversely, the model trained on the proposed-preprocessed data was tested on baseline-preprocessed data. This setup allows us to evaluate whether the learned representations remain stable under distribution shifts caused by different preprocessing strategies [23]. It also helps determine whether the model relies on preprocessing-specific characteristics or captures more generalizable neural patterns.

To investigate spectral dependence, frequency ablation experiments were performed by selectively removing specific EEG frequency bands (delta, theta, alpha, beta, and gamma), including the line-noise band (45–55 Hz), prior to model training. Temporal ablation analysis was also conducted by restricting the input to specific post-stimulus time windows to evaluate the contribution of temporally localized neural components, particularly those associated with visual ERP responses in the RSVP paradigm [24].

In addition, robustness to signal perturbation was assessed by injecting Gaussian noise with varying standard deviations (*σ* = 0.0, 0.1, 0.3, 0.5, 1.0) during testing. The resulting degradation in classification accuracy was analyzed to evaluate model stability under increasing noise levels [25].

### E. Effective Connectivity Matrices

In this study, PDC, DTF, and GPDC are treated as biologically motivated reference measures of effective connectivity.Since they provide model-based estimates of directed neural interactions, their role in this study is to serve as principled reference representations against which the learned attention matrices can be compared [7]. This process is generally based on an MVAR model that describes the interaction between signals *X*(*t*) as a linear combination of their past values:

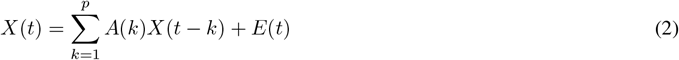

where *A*(*k*) is model coefficient matrix for every *lag k* and *E*(*t*) is the white noise. To ensure an optimal balance between model complexity and goodness-of-fit, the MVAR model order *p* was determined using the Bayesian Information Criterion (BIC). We evaluated candidate orders ranging from 5 to 20 across representative trial samples. Based on the median of the optimal BIC scores, the model order was systematically established at *p* = 5. The frequency-domain connectivity metrics were subsequently derived using a 512-point Fast Fourier Transform (FFT).

Based on this model, PDC is used to observe only direct causal interactions between two processes in a multivariate system by normalizing the elements of the coefficient matrix in the frequency domain. PDC can be formulated as:

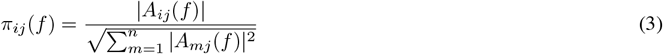

GPDC is a further development of PDC with scale-invariance properties that can overcome the weakness of PDC in terms of amplitude differences between signals. GPDC can be formulated as:

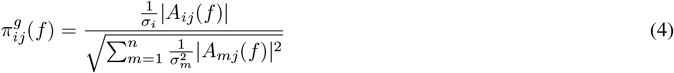

On the other hand, DTF measures the total causal influence, both direct and indirect, from one channel to another. DTF measurement uses the ratio of the transfer matrix element to the total influence received. DTF can be formulated as:

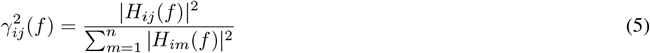

### F. Attention-Connectivity Comparative Analysis

To assess the biological plausibility of the learned attention representations, the channel-wise attention weight matrices were compared with the effective connectivity matrices constructed using PDC, GPDC, and DTF. The comparative flow can be seen in Fig. 1B. For this analysis, the attention matrices were extracted from the model at the final training epoch (epoch 40) and averaged across all 4 attention heads, as well as across all test trials for all 10 subjects, to obtain a stable global attention representation. The effective connectivity matrices were computed subject-wise and trial-wise using the same test data, and then averaged across trials and subjects to obtain stable reference connectivity representations for each frequency band and connectivity metric.

We evaluated the alignment between Transformer attention and EEG connectivity using two complementary metrics: node-strength correlation and representational similarity analysis (RSA). Node-strength correlation focuses on local topology, testing whether the most active channels (hubs) in the attention graph match those in the EEG connectivity graph. Conversely, RSA evaluates the global network structure, measuring whether the overall pattern of pairwise relationships is similar across both representations. Because node-strength targets local hubs while RSA targets global patterns, a divergence in their results is not contradictory. This divergence suggests that preprocessing may selectively influence the scale at which biological alignment is preserved, favoring global representational geometry over local hub structure.

Technically, for the node-strength correlation, total node strength was computed by summing the weighted incoming and outgoing connections per channel. Spearman rank correlation was then calculated between the node strengths derived from the attention matrices and those derived from the effective connectivity matrices across the evaluated frequency bands [26]. Furthermore, RSA was performed by constructing representational dissimilarity matrices (RDMs) from both the attention and connectivity matrices. The similarity between these RDMs was quantified using Spearman rank correlation to evaluate the extent of representational correspondence between the learned attention patterns and the directed effective connectivity structures [27].

### G. Statistical Analysis

To evaluate the statistical significance of performance differences and structural alignments across all experimental conditions, we systematically conducted paired t-tests. The evaluated conditions encompassed the comparison of preprocessing pipelines, frequency and temporal ablations, noise injection robustness, node-strength correlations, and representational similarity analysis (RSA) scores. A paired design was strictly chosen because all evaluations were performed on the exact same set of 10 subjects [28]. For the ablation studies specifically, statistical significance was calculated by comparing the performance of each ablated condition against the un-ablated reference model (denoted as ‘None’).

To quantify the magnitude of these differences beyond mere statistical significance, effect sizes were calculated using Cohen’s *d* for paired samples. The effect size was defined as the mean of the paired differences divided by the sample standard deviation of the differences [29].

All classification performance metrics (Top-1 and Top-5 accuracies) are reported as percentages, and together with the comparative graph metrics, are presented in the format of mean *±* standard deviation (SD) across the subjects.

To rigorously control for the multiple comparison problem inherent in evaluating numerous conditions, we applied the False Discovery Rate (FDR) correction using the Benjamini-Hochberg procedure. The FDR correction was applied independently within each distinct type of analyses to preserve statistical power while strictly controlling the expected proportion of false positives [30]. Throughout the results, statistical significance is explicitly denoted based on these FDR-corrected *p*-values (*p*_*FDR*_) at two sequential thresholds: *p*_*FDR*_ *<* 0.05 (*) and *p*_*FDR*_ *<* 0.01 (**).

### IV. Results and Discussion

### A. Effect of Preprocessing on Spectral Characteristics

The performance of deep learning models is highly dependent on the input data. EEG signals naturally contain a lot of noise or interference, so effective preprocessing is required before evaluating the accuracy of the model [31]. Therefore, we analyzed the power spectral density (PSD) in the 1-80 Hz frequency range in the frontal, occipital, and whole head areas (Fig. 2). This was done to see how cleaning components using ICA, filters, and MVNN changed the signal power.

**Fig. 2.**
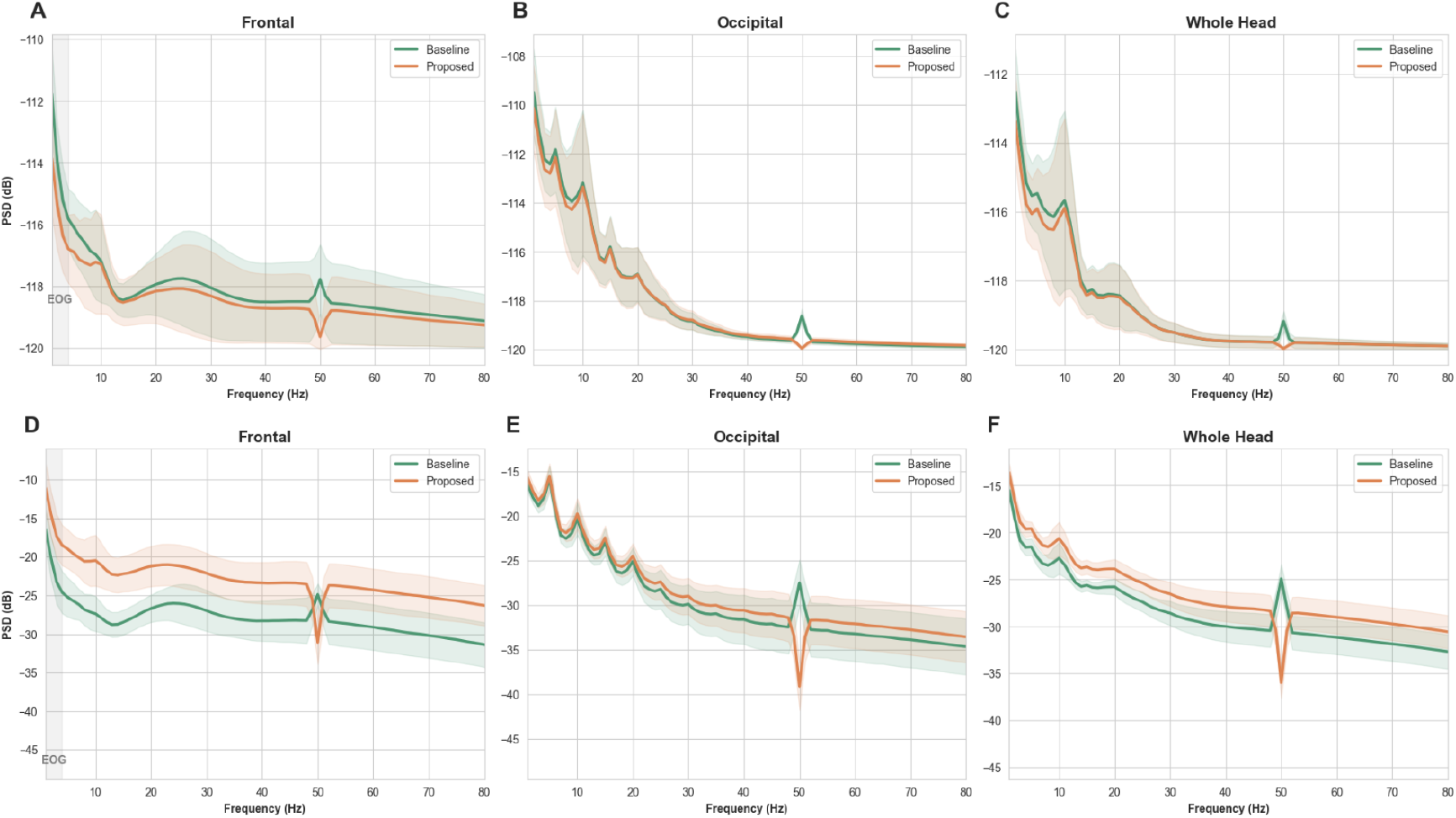
Power Spectral Density (PSD) analysis comparing the baseline and proposed preprocessing pipelines across different brain regions. (A)-(C) PSD distribution in the Frontal, Occipital, and Whole Head regions before MVNN normalization, highlighting the suppression of line noise (50 Hz) and frontal artifacts in the proposed pipeline. (D)-(F) The corresponding PSD distributions after MVNN normalization, demonstrating a more balanced spectral power across regions.

In the PSD visualization before MVNN, the proposed preprocessing pipeline shows a clear reduction of spectral power in the frontal region compared to the baseline condition. This decrease is consistent with the effect of ICA, which is commonly used to remove eye blink artifacts that affect the frontal electrodes [17]. Moreover, a clear drop around 50 Hz can be observed in the proposed condition. This indicates that the applied notch filter effectively suppresses line noise that is still visible in the baseline signal [32].

After MVNN normalization, the proposed condition showed relatively higher and more evenly distributed power across scalp regions. This pattern likely reflects the prior removal of non-neural artifacts and the suppression of certain frequency disturbances which results in neural activity becoming more prominent in the normalized signal [33].

### B. Impact of Preprocessing on Model Performance

The learning dynamics during 40 epochs are shown in Fig. 3A and Fig. 3B, which displays the Top-1 and Top-5 accuracy curves throughout the training process. The curves of both models show a stable convergence pattern without extreme fluctuations, indicating that the optimization process proceeded consistently under both preprocessing conditions. No significant overfitting was observed towards the end of the epoch, so the use of the model at epoch 40 as the final representation is considered sufficiently representative [34].

**Fig. 3.**
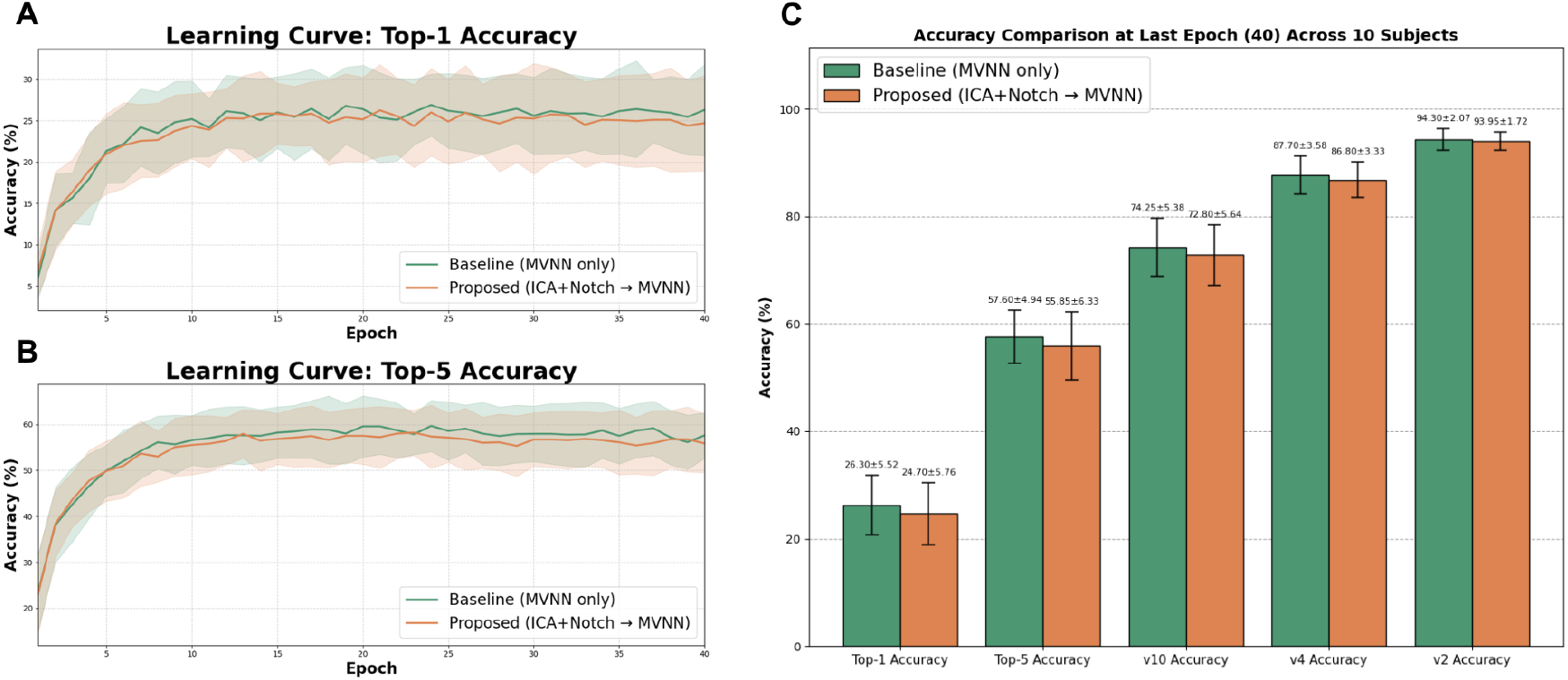
Model training dynamics and classification performance. (A) Top-1 and (B) Top-5 accuracy learning curves over 40 epochs, showing stable convergence for both baseline and proposed models. (C) Comprehensive accuracy comparison (Top-1, Top-5, V10, V4, and V2 settings) evaluated at the final epoch across 10 subjects.

The main accuracy evaluation results are presented in Fig. 3C, which summarizes the average performance across subjects for the Top-1, Top-5, and V2, V4, and V10 metrics. In general, the model with baseline preprocessing shows slightly higher performance than the proposed model. The average Top-1 accuracy on the baseline reached 26.30% ± 5.52, while the proposed model achieved 24.70% ± 5.76. A similar pattern was also seen in the Top-5 accuracy (57.60% vs. 55.85%), as well as in the V10, V4, and V2 scenarios, where the baseline consistently outperformed by a relatively small margin. However, the differences between conditions were still within a close standard deviation range, indicating that both pipelines produced generally comparable levels of performance [35].

To evaluate cross-preprocessing generalization, a cross-test analysis was conducted and summarized in Table I. The average accuracy shows that the baseline model tested on the proposed data has identical average performance (0.263), while the proposed model tested on the baseline data produces an average of 0.241. When tested on the pipeline corresponding to its training, the baseline achieved 0.263 and the proposed model achieved 0.247. In general, there was no drastic decrease in performance due to the exchange of the test pipeline, although there were variations between subjects. This indicates that the representation learned by the model does not depend entirely on the specific characteristics of preprocessing, but there are still indications of a shift in distribution that affects performance stability in certain subjects [21], [22].

**TABLE I.** TOP-1 ACCURACY COMPARISON BETWEEN BASELINE AND PROPOSED PREPROCESSING PIPELINES ACROSS SUBJECTS.

| Sub | Baseline Training |  | Proposed Training |  |
| --- | --- | --- | --- | --- |
|  | Test-B | Test-P | Test-B | Test-P |
| 1 | 0.235 | 0.240 | 0.245 | 0.245 |
| 2 | 0.225 | 0.255 | 0.210 | 0.200 |
| 3 | 0.235 | 0.240 | 0.255 | 0.255 |
| 4 | 0.225 | 0.250 | 0.220 | 0.190 |
| 5 | 0.210 | 0.210 | 0.200 | 0.175 |
| 6 | 0.265 | 0.240 | 0.270 | 0.245 |
| 7 | 0.260 | 0.275 | 0.200 | 0.230 |
| 8 | 0.395 | 0.380 | 0.355 | 0.360 |
| 9 | 0.265 | 0.230 | 0.135 | 0.245 |
| 10 | 0.315 | 0.310 | 0.315 | 0.325 |
| M | $0.263 \pm 0.06$ | $0.263 \pm 0.05$ | $0.241 \pm 0.06$ | $0.247 \pm 0.06$ |

Overall, these results are consistent with our first hypothesis, indicating that the proposed pipeline achieves more comprehensive signal cleaning without substantially compromising decoding performance. Although the artifact removal process did not lead to a direct increase in classification accuracy, the performance of the two approaches remained relatively comparable, with the baseline only outperforming the proposed model by a small, non-significant margin. This suggests that the model is robust to the distribution shifts introduced by rigorous cleaning and that the effect of preprocessing on generalizability is moderate compared to the dominance of the core representation factors learned during the training process. These findings provide a critical foundation for our subsequent analyses, as they confirm that the observed attention patterns are derived from a cleaner neural signal while maintaining stable predictive power.

### C. Spectral Ablation

The results, detailed in Table II and visualized in Fig. 4A and 4B, demonstrate the distinct contribution of each frequency band to the model’s predictive performance. In general, the removal of low frequencies causes the most substantial decrease in performance across both models. Ablating the theta (*θ*) band drastically reduces Top-1 accuracy from 26.30% to 7.50% in the baseline model and from 24.70% to 8.10% in the proposed model (*p <* 0.001). This decline is closely followed by the removal of the delta (*δ*) band, which significantly reduces accuracy to approximately 11–12% (*p <* 0.001).

**TABLE II.** Spectral Ablation Analysis Results.

| Band | Top-1 Acc (%) |  |  | Top-5 Acc (%) |  |  |
| --- | --- | --- | --- | --- | --- | --- |
| | Acc ± SD | *p*-value | *d* | Acc ± SD | *p*-value | *d* |
| **Baseline Model** |  |  |  |  |  |  |
| None | 26.30 ± 5.52 | - | - | 57.65 ± 4.86 | - | - |
| LN | 26.35 ± 5.48 | 0.591 | 0.18 | 57.25 ± 5.08 | 0.167 | -0.49 |
| $\gamma$ | 25.35 ± 5.07 | 0.084 | -0.65 | 56.45 ± 5.34 | 0.063 | -0.73 |
| $\beta$ | 18.55 ± 5.19 | <0.001** | -1.83 | 45.05 ± 7.87 | <0.001** | -1.90 |
| $\alpha$ | 21.85 ± 5.38 | 0.045* | -0.81 | 49.45 ± 8.61 | 0.003** | -1.41 |
| $\theta$ | 7.50 ± 4.56 | <0.001** | -3.71 | 22.90 ± 11.08 | <0.001** | -3.28 |
| $\delta$ | 11.95 ± 2.03 | <0.001** | -3.11 | 33.00 ± 5.65 | <0.001** | -5.47 |
| **Proposed Model** |  |  |  |  |  |  |
| None | 24.70 ± 5.76 | - | - | 55.85 ± 6.33 | - | - |
| LN | 24.90 ± 5.96 | 0.323 | 0.37 | 55.80 ± 6.26 | 0.811 | -0.08 |
| $\gamma$ | 24.40 ± 5.94 | 0.390 | -0.31 | 55.25 ± 6.50 | 0.058 | -0.74 |
| $\beta$ | 18.50 ± 5.88 | 0.007** | -1.19 | 44.85 ± 9.40 | 0.002** | -1.48 |
| $\alpha$ | 19.70 ± 6.12 | 0.002** | -1.62 | 48.00 ± 9.11 | 0.003** | -1.37 |
| $\theta$ | 8.10 ± 5.03 | <0.001** | -2.56 | 24.20 ± 12.33 | <0.001** | -2.73 |
| $\delta$ | 11.85 ± 2.76 | <0.001** | -3.10 | 33.95 ± 6.08 | <0.001** | -4.20 |

\* $p < 0.05$ , \*\* $p < 0.01$ (FDR-corrected, compared to None).
*d* represents Cohen's *d*. LN = Line Noise (45-55 Hz).

**Fig. 4.**
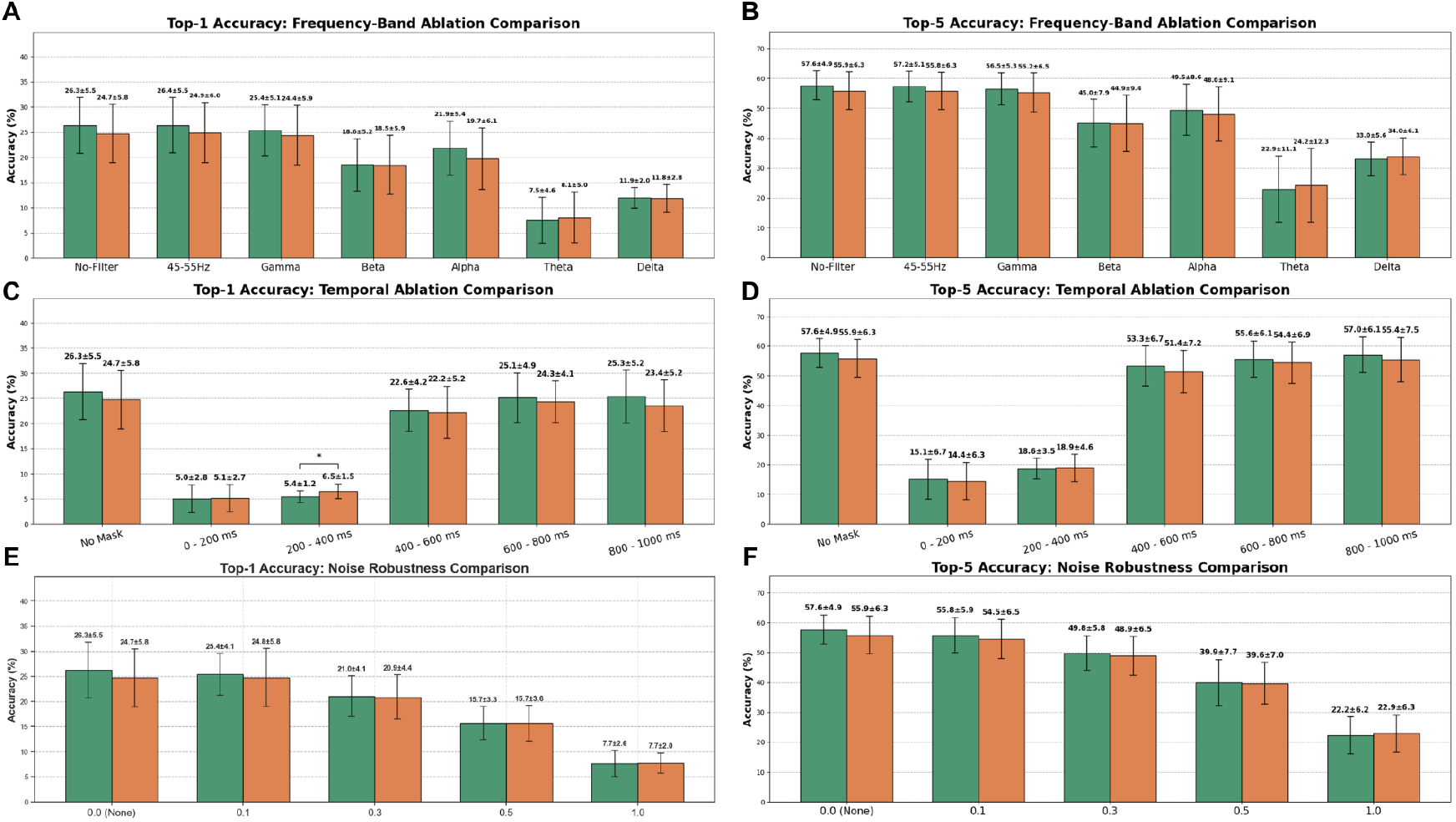
Ablation study results evaluating the contribution of specific EEG components to model performance. (A) Top-1 and (B) Top-5 accuracy drops following the removal of specific frequency bands, indicating the critical role of low frequencies (*θ* and *δ*). (C) Top-1 and (D) Top-5 accuracy drops following the removal of specific time windows, highlighting the importance of early visual ERP responses (0–400 ms). (E) Top-1 and (F) Top-5 accuracy show a consistent and significant decline in performance for both models as the noise intensity levels increase from 0.0 to 1.0.

These steep declines are consistently observed in both Top-1 and Top-5 accuracy metrics, confirming that essential visual decoding information in the RSVP dataset is heavily concentrated in the low-frequency spectrum. This finding is consistent with established neurophysiological principles, as visual processing in rapid paradigms is commonly associated with strong delta and theta-band ERP components [36]–[38].

Furthermore, the removal of the alpha (*α*) and beta (*β*) bands also reduces performance, although with a more moderate impact that brings Top-1 accuracies down to the 18–22% range. Conversely, filtering out the gamma (*γ*) band and line noise (LN, 45–55 Hz) components exerts no statistically significant negative effect (*p >* 0.05) on the models’ predictive capabilities.

The nearly identical ablation degradation patterns observed between the baseline and proposed models suggest that both preprocessing pipelines successfully preserve similar fundamental spectral information structures, regardless of the additional artifact removal steps in the proposed pipeline [38]. This stability in the underlying feature importance provides a clear transition to evaluating whether these spectral components are also localized within specific time windows.

### D. Temporal Ablation

As detailed in the results (Table III, Fig. 4C, and Fig. 4D), the removal of activity within the 0 to 200 ms window (*T*_1_) caused a substantial decrease in performance. Top-1 accuracy plummeted from 26.30% to 5.00% in the baseline model and from 24.70% to 5.10% in the proposed model (*p <* 0.001). A similarly steep decline was observed in the Top-5 metric, indicating that the early phase of neural responses plays a critical role in capturing the initial perceptual processing of the visual stimulus [39]. Crucially, this 200 ms time frame corresponds exactly to the duration of a single trial in our RSVP paradigm, which consists of a 100 ms image presentation followed by a 100 ms blank inter-stimulus interval [20]. Therefore, ablating this specific window essentially eliminates the primary evoked potentials generated during the exact physical presence and immediate afterimage of the stimulus.

**TABLE III.** Temporal Ablation Analysis Results.

| TW | Top-1 Acc (%) |  |  | Top-5 Acc (%) |  |  |
| --- | --- | --- | --- | --- | --- | --- |
| | Acc ± SD | $p$-value | $d$ | Acc ± SD | $p$-value | $d$ |
| **Baseline Model** |  |  |  |  |  |  |
| None | 26.30 ± 5.52 | - | - | 57.65 ± 4.86 | - | - |
| $T_1$ | 5.00 ± 2.77 | <0.001** | -4.00 | 15.10 ± 6.65 | <0.001** | -6.85 |
| $T_2$ | 5.40 ± 1.17 | <0.001** | -4.35 | 18.65 ± 3.54 | <0.001** | -7.15 |
| $T_3$ | 22.55 ± 4.17 | <0.001** | -2.03 | 53.30 ± 6.70 | 0.008** | -1.17 |
| $T_4$ | 25.10 ± 4.90 | 0.055 | -0.73 | 55.60 ± 6.12 | 0.030* | -0.88 |
| $T_5$ | 25.30 ± 5.24 | 0.055 | -0.72 | 57.00 ± 6.06 | 0.380 | -0.29 |
| **Proposed Model** |  |  |  |  |  |  |
| None | 24.70 ± 5.76 | - | - | 55.85 ± 6.33 | - | - |
| $T_1$ | 5.10 ± 2.70 | <0.001** | -3.75 | 14.45 ± 6.32 | <0.001** | -5.39 |
| $T_2$ | 6.50 ± 1.45 | <0.001** | -3.48 | 18.90 ± 4.56 | <0.001** | -6.18 |
| $T_3$ | 22.20 ± 5.18 | 0.012* | -1.09 | 51.40 ± 7.16 | 0.002** | -1.49 |
| $T_4$ | 24.30 ± 4.10 | 0.542 | -0.20 | 54.40 ± 6.94 | 0.026* | -0.88 |
| $T_5$ | 23.45 ± 5.15 | 0.012* | -1.06 | 55.40 ± 7.52 | 0.500 | -0.25 |

\* $p < 0.05$ , \*\* $p < 0.01$ (FDR-corrected, compared to None).
$d$ represents Cohen's $d$ . TW = Time Window (ms).
$T_1$ (0–200), $T_2$ (200–400), $T_3$ (400–600), $T_4$ (600–800), $T_5$ (800–1000).

An even stronger effect size was observed when the 200 to 400 ms time window (*T*_2_) was removed, a period closely associated with the P300 ERP component. Under this condition, Top-1 accuracy dropped to approximately 5 to 6% (*p <* 0.001) in both models, accompanied by the largest Cohen’s *d* values recorded in this study (up to *−*4.35 for Top-1 and *−*7.15 for Top-5). In contrast, ablating activity in the later windows from 400 to 1000 ms (*T*_3_ through *T*_5_) caused only moderate or statistically insignificant decreases in performance.

These results support our second hypothesis, indicating that the most essential information for visual decoding is primarily localized within the first 400 ms after stimulus onset. This temporal pattern strongly aligned with recent deep learning-based EEG studies, which indicate that visual representations are largely formed within this early window, after which decoding performance tends to stabilize around 500 ms [2], [36]. Having identified these internal spectral and temporal drivers of classification, we naturally progress to evaluating how resilient these learned features are against external signal interference in the noise robustness analysis.

### E. Noise Robustness

Having established the internal temporal and spectral dependencies of the model, we next evaluated its resilience against external signal interference. The noise robustness experiment tests how well both preprocessing pipelines maintain their predictive performance when subjected to increasing levels of Gaussian noise (Table IV, Fig. 4E, and Fig. 4F). As detailed in Table IV, statistical significance was calculated by comparing the performance at each noise intensity against the noise-free reference condition (*σ* = 0.0).

**TABLE IV.** Noise Robustness Analysis Results.

| $\sigma$ | Top-1 Acc (%) | | | Top-5 Acc (%) | | |
| --- | --- | --- | --- | --- | --- | --- |
| | Acc ± SD | $p$-value | $d$ | Acc ± SD | $p$-value | $d$ |
| **Baseline Model** |  |  |  |  |  |  |
| 0.0 | 26.30 ± 5.52 | - | - | 57.65 ± 4.86 | - | - |
| 0.1 | 25.40 ± 4.14 | 0.260 | -0.38 | 55.75 ± 5.93 | 0.007** | -1.13 |
| 0.3 | 21.00 ± 4.07 | <0.001** | -1.80 | 49.80 ± 5.76 | <0.001** | -2.24 |
| 0.5 | 15.70 ± 3.30 | <0.001** | -2.86 | 39.90 ± 7.67 | <0.001** | -5.34 |
| 1.0 | 7.65 ± 2.61 | <0.001** | -5.46 | 22.20 ± 6.21 | <0.001** | -8.12 |
| **Proposed Model** |  |  |  |  |  |  |
| 0.0 | 24.70 ± 5.76 | - | - | 55.85 ± 6.33 | - | - |
| 0.1 | 24.75 ± 5.76 | 0.864 | 0.06 | 54.50 ± 6.52 | 0.004** | -1.25 |
| 0.3 | 20.85 ± 4.42 | 0.002** | -1.47 | 48.85 ± 6.49 | <0.001** | -3.06 |
| 0.5 | 15.65 ± 3.56 | <0.001** | -2.48 | 39.65 ± 6.97 | <0.001** | -4.01 |
| 1.0 | 7.70 ± 2.02 | <0.001** | -3.62 | 22.85 ± 6.34 | <0.001** | -5.04 |

\* $p < 0.05$ , \*\* $p < 0.01$ (FDR-corrected, compared to $\sigma = 0.0$ ).
$d$ represents Cohen's $d$ . $\sigma$ indicates the standard deviation of injected Gaussian noise.

At a low noise level (*σ* = 0.1), Top-1 accuracy remains statistically stable (*p >* 0.05 for both models), although the Top-5 metric begins to show a slight but significant decrease. When the noise level increases to *σ* = 0.3, Top-1 accuracy experiences a significant drop to 21.00% in the baseline model and 20.85% in the proposed model (*p <* 0.01).

The performance degradation becomes substantially more pronounced at *σ* = 0.5, with Top-1 accuracy falling to the 15 to 16% range (*p <* 0.001). At the highest noise intensity (*σ* = 1.0), accuracy plummets to approximately 7 to 8%, accompanied by extremely large effect sizes (Cohen’s *d* reaching *−*5.46 for the baseline and *−*3.62 for the proposed model).

The nearly identical performance degradation trajectories between the two models indicate that both preprocessing pipelines possess comparable levels of noise robustness [25]. This confirms that the rigorous artifact removal in the proposed pipeline does not make the model more vulnerable to random signal fluctuations. With the technical stability and feature dependencies of the models thoroughly confirmed, we can now shift our focus to evaluating their biological interpretability through network alignment analyses.

### F. Alignment Between Model Attention and Effective Connectivity

Node strength analysis was first used to evaluate how the learned representations relate to local topological hubs in the brain network during visual processing. As shown in Table V and Fig. 5A, the correlation values in the proposed preprocessing pipeline generally exhibit a nominal decrease compared to the baseline, particularly in the DTF and PDC metrics across several frequency bands. These nominal decreases are accompanied by moderate to large effect sizes (for example, Cohen’s *d* ranging from *−*0.7 to *−*1.03 in the alpha and beta bands, which are typically associated with long-range communication and top-down control mechanisms [40], [41]). However, after rigorous FDR correction, none of these differences reach statistical significance (all *p >* 0.05). This indicates that, although the proposed preprocessing pipeline showed a nominal decrease in several node-strength correlations, these differences did not remain statistically significant after FDR correction.

**Fig. 5.**
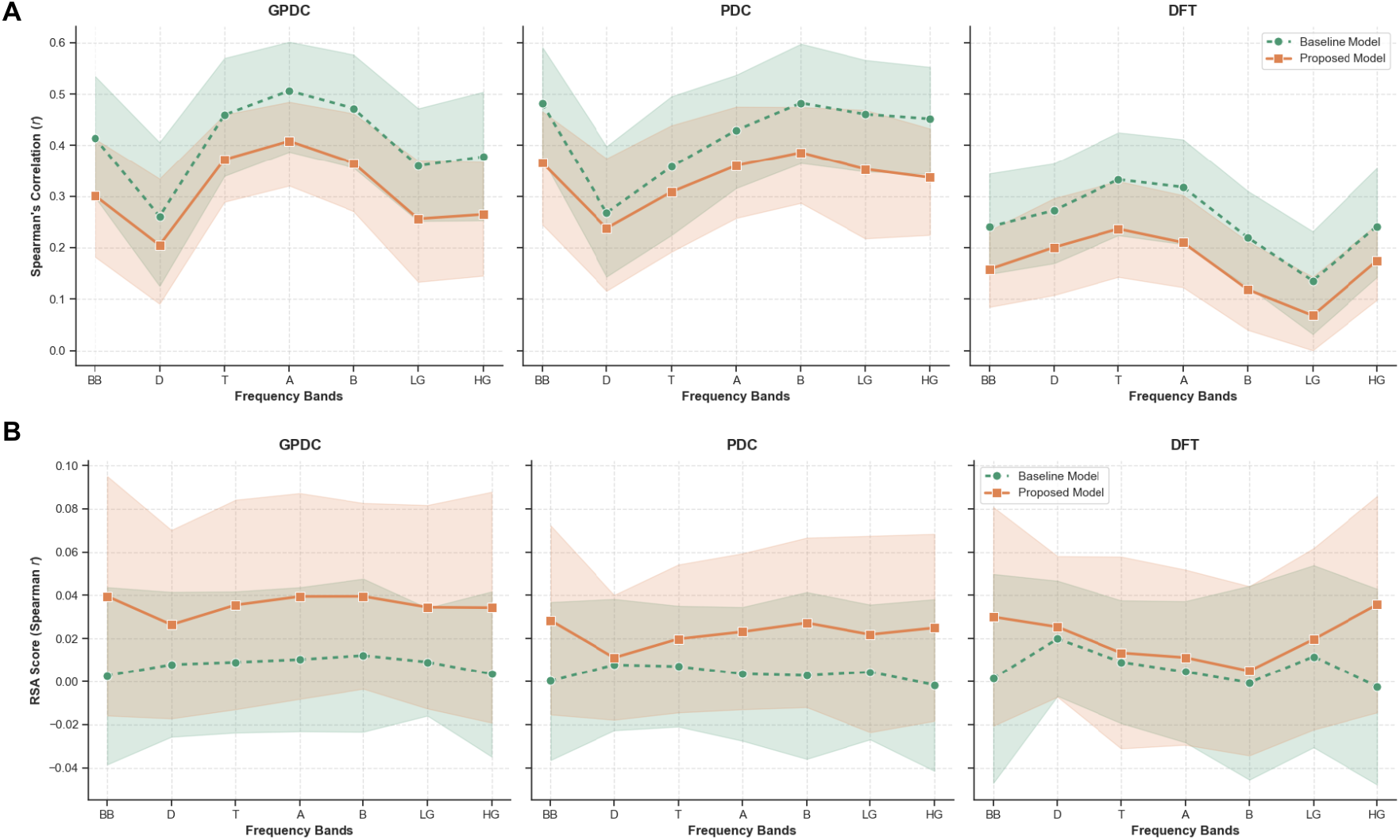
Alignment analysis between the data-driven AI attention weights and the biological effective connectivity (GPDC, PDC, and DTF) across broadband (BB) and specific frequency bands. (A) Spearman’s correlation of node strength, comparing network topology between the models. (B) Representational Similarity Analysis (RSA) scores, showing a general trend toward higher RSA-based structural alignment under the proposed preprocessing pipeline.

**TABLE V.** Node Strength Correlation Analysis.

| Method | Band | Incoming Strength |  |  |  |  | Outgoing Strength |  |  |  |  | Total Node Strength |  |  |  |  |
| --- | --- | --- | --- | --- | --- | --- | --- | --- | --- | --- | --- | --- | --- | --- | --- | --- |
|  |  | Base | Prop | Diff | p-value | d | Base | Prop | Diff | p-value | d | Base | Prop | Diff | p-value | d |
| DTF | BB | 0.065 | 0.003 | -0.062 | 0.127 | -0.87 | -0.253 | -0.343 | -0.090 | 0.127 | -0.80 | 0.241 | 0.158 | -0.083 | 0.127 | -0.69 |
| | $\delta$ | 0.187 | 0.122 | -0.065 | 0.127 | -0.76 | -0.139 | -0.222 | -0.083 | 0.127 | -0.80 | 0.272 | 0.201 | -0.072 | 0.226 | -0.48 |
| | $\theta$ | 0.275 | 0.214 | -0.061 | 0.140 | -0.64 | -0.105 | -0.186 | -0.082 | 0.127 | -0.77 | 0.333 | 0.237 | -0.096 | 0.142 | -0.62 |
| | $\alpha$ | 0.262 | 0.193 | -0.069 | 0.127 | -0.80 | -0.119 | -0.202 | -0.083 | 0.127 | -0.72 | 0.318 | 0.210 | -0.108 | 0.127 | -0.74 |
| | $\beta$ | 0.113 | 0.045 | -0.068 | 0.127 | -1.03 | -0.237 | -0.341 | -0.104 | 0.127 | -0.77 | 0.220 | 0.118 | -0.101 | 0.127 | -0.92 |
| | Low $\gamma$ | -0.046 | -0.097 | -0.051 | 0.127 | -0.75 | -0.333 | -0.426 | -0.093 | 0.127 | -0.68 | 0.136 | 0.068 | -0.068 | 0.140 | -0.64 |
| | High $\gamma$ | 0.015 | -0.037 | -0.053 | 0.127 | -0.68 | -0.241 | -0.314 | -0.074 | 0.127 | -0.69 | 0.241 | 0.174 | -0.067 | 0.152 | -0.60 |
| GPDC | BB | 0.342 | 0.297 | -0.045 | 0.361 | -0.34 | -0.468 | -0.535 | -0.067 | 0.295 | -0.40 | 0.415 | 0.302 | -0.113 | 0.137 | -0.66 |
| | $\delta$ | 0.461 | 0.428 | -0.033 | 0.607 | -0.18 | 0.213 | 0.199 | -0.015 | 0.727 | -0.11 | 0.261 | 0.205 | -0.056 | 0.195 | -0.51 |
| | $\theta$ | 0.538 | 0.508 | -0.030 | 0.607 | -0.17 | 0.094 | 0.044 | -0.050 | 0.246 | -0.46 | 0.459 | 0.372 | -0.087 | 0.195 | -0.53 |
| | $\alpha$ | 0.541 | 0.510 | -0.031 | 0.573 | -0.20 | -0.114 | -0.166 | -0.052 | 0.195 | -0.52 | 0.506 | 0.408 | -0.098 | 0.195 | -0.53 |
| | $\beta$ | 0.449 | 0.410 | -0.039 | 0.442 | -0.28 | -0.423 | -0.484 | -0.060 | 0.259 | -0.44 | 0.472 | 0.365 | -0.107 | 0.195 | -0.52 |
| | Low $\gamma$ | 0.269 | 0.231 | -0.038 | 0.400 | -0.31 | -0.465 | -0.519 | -0.054 | 0.336 | -0.36 | 0.360 | 0.256 | -0.104 | 0.168 | -0.58 |
| | High $\gamma$ | 0.286 | 0.240 | -0.046 | 0.336 | -0.36 | -0.452 | -0.516 | -0.064 | 0.336 | -0.36 | 0.378 | 0.265 | -0.113 | 0.127 | -0.73 |
| PDC | BB | 0.049 | -0.009 | -0.059 | 0.127 | -0.79 | -0.268 | -0.355 | -0.087 | 0.127 | -0.81 | 0.481 | 0.365 | -0.116 | 0.141 | -0.63 |
| | $\delta$ | 0.143 | 0.079 | -0.064 | 0.127 | -0.89 | -0.079 | -0.145 | -0.067 | 0.169 | -0.57 | 0.268 | 0.238 | -0.030 | 0.504 | -0.24 |
| | $\theta$ | 0.206 | 0.157 | -0.049 | 0.127 | -0.69 | -0.110 | -0.181 | -0.070 | 0.142 | -0.62 | 0.358 | 0.308 | -0.049 | 0.405 | -0.30 |
| | $\alpha$ | 0.222 | 0.173 | -0.049 | 0.127 | -0.69 | -0.145 | -0.222 | -0.077 | 0.127 | -0.69 | 0.429 | 0.360 | -0.069 | 0.336 | -0.37 |
| | $\beta$ | 0.154 | 0.100 | -0.054 | 0.127 | -0.78 | -0.214 | -0.294 | -0.080 | 0.127 | -0.71 | 0.483 | 0.386 | -0.097 | 0.262 | -0.43 |
| | Low $\gamma$ | 0.029 | -0.024 | -0.053 | 0.127 | -0.72 | -0.283 | -0.361 | -0.078 | 0.127 | -0.72 | 0.461 | 0.353 | -0.108 | 0.195 | -0.52 |
| | High $\gamma$ | 0.008 | -0.045 | -0.053 | 0.127 | -0.70 | -0.304 | -0.396 | -0.092 | 0.127 | -0.85 | 0.452 | 0.337 | -0.115 | 0.127 | -0.76 |

\* $p < 0.05$ , \*\* $p < 0.01$ . Base = Baseline, Prop = Proposed, Diff = Mean Difference (Prop-Base), $d$ = Cohen's $d$ . BB = Broadband.

**TABLE VI.** Representational Similarity Analysis (RSA) Results.

| Method | Band | Base | Prop | Diff | *p*-value | *d* |
| --- | --- | --- | --- | --- | --- | --- |
| DTF | BB | 0.001 | 0.030 | 0.029 | 0.356 | 0.49 |
| | $\delta$ | 0.020 | 0.025 | 0.006 | 0.719 | 0.20 |
| | $\theta$ | 0.009 | 0.013 | 0.004 | 0.777 | 0.09 |
| | $\alpha$ | 0.005 | 0.011 | 0.007 | 0.777 | 0.14 |
| | $\beta$ | -0.001 | 0.005 | 0.006 | 0.777 | 0.10 |
| | Low $\gamma$ | 0.012 | 0.020 | 0.008 | 0.777 | 0.12 |
| | High $\gamma$ | -0.002 | 0.036 | 0.038 | 0.356 | 0.62 |
| GPDC | BB | 0.003 | 0.040 | 0.037 | 0.356 | 0.66 |
| | $\delta$ | 0.008 | 0.027 | 0.019 | 0.458 | 0.35 |
| | $\theta$ | 0.009 | 0.036 | 0.027 | 0.356 | 0.44 |
| | $\alpha$ | 0.010 | 0.040 | 0.029 | 0.356 | 0.48 |
| | $\beta$ | 0.012 | 0.040 | 0.027 | 0.356 | 0.49 |
| | Low $\gamma$ | 0.009 | 0.035 | 0.025 | 0.356 | 0.45 |
| | High $\gamma$ | 0.003 | 0.034 | 0.031 | 0.356 | 0.56 |
| PDC | BB | 0.000 | 0.028 | 0.028 | 0.356 | 0.56 |
| | $\delta$ | 0.008 | 0.011 | 0.003 | 0.777 | 0.10 |
| | $\theta$ | 0.007 | 0.020 | 0.013 | 0.458 | 0.34 |
| | $\alpha$ | 0.004 | 0.023 | 0.020 | 0.356 | 0.43 |
| | $\beta$ | 0.003 | 0.027 | 0.025 | 0.356 | 0.48 |
| | Low $\gamma$ | 0.004 | 0.022 | 0.018 | 0.464 | 0.33 |
| | High $\gamma$ | -0.002 | 0.025 | 0.027 | 0.356 | 0.48 |

Base = Mean Baseline, Prop = Mean Proposed, Diff = Mean Difference.
*d* represents Cohen's *d*. BB = Broadband.

Furthermore, the GPDC method demonstrated even smaller mean differences, reaffirming that the global directional connectivity structure remains highly stable [42]. Because no statistically significant differences were found across any frequency bands or connectivity metrics, the overall hub-level alignment of the network appears broadly preserved across preprocessing strategies [43]. Thus, the models maintain a comparable degree of alignment with established node centrality measures regardless of the applied preprocessing techniques. This suggests that preprocessing differences likely affected only subtle local variances rather than the global organization of the network during visual decoding.

Beyond the comparative stability between the pipelines, the absolute magnitude of these correlations provides critical insights into the biological validity of the models. Across all evaluated metrics, GPDC demonstrated the highest structural alignment with the data-driven attention, yielding moderate to strong positive correlations (ranging from 0.40 to 0.54 for total node strength), particularly in the *θ* and *α* bands. In contrast, alignments evaluated using DTF and PDC yielded notably weaker correlations that were often near zero. This substantial difference indicates that the AI’s learned attention best captures the specific connectivity dynamics modeled by GPDC.

To further illustrate these structural characteristics, Fig. 6 visualizes the spatial mapping by comparing the spatial patterns of the AI’s channel-wise attention against the reference EEG effective connectivity. Based on the absolute alignment strengths, GPDC was naturally selected for this visualization. Specifically, the *θ* and *α* bands were chosen because the spectral ablation study identified *θ* as the most critical frequency band for predictive accuracy, while the *α* band was considered due to its established role in visual attention and top-down cortical processing [41]. The topographical maps of the incoming, outgoing, and total node strengths reveal a distinct spatial alignment between the data-driven attention weights and the neurophysiological reference networks. Notably, the attention matrices from both the baseline and proposed models highlight regions that are structurally consistent with the brain’s established connectivity hubs during visual processing [36], [44]. Furthermore, the visual comparison between the two preprocessing pipelines shows highly similar topographical distributions. This qualitative visualization is consistent with the preceding statistical findings, suggesting that the broad spatial organization of the learned attention patterns remains similar across preprocessing strategies [45].

**Fig. 6.**
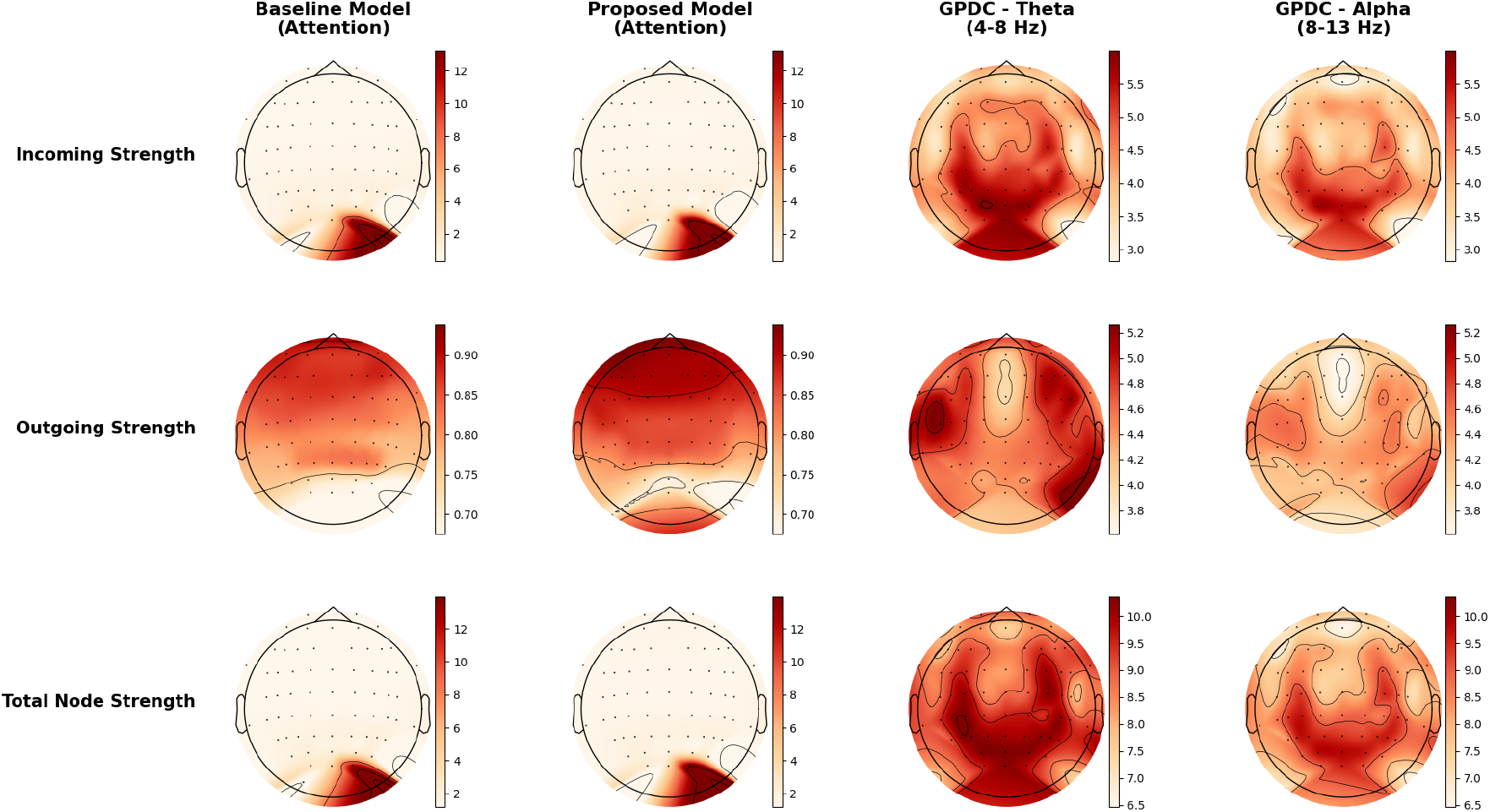
Topographical comparison between AI channel-wise attention and reference EEG effective connectivity. These maps illustrate the spatial patterns of Incoming, Outgoing, and Total Node Strengths across 63 EEG channels. The first two columns display the data-driven attention weights extracted from the baseline and proposed models. The last two columns display the corresponding reference connectivity derived from the GPDC metric in the *θ* (4–8 Hz) and *α* (8–13 Hz) frequency bands. Color scales are normalized independently for the attention and connectivity representations to emphasize the alignment of spatial network patterns rather than absolute magnitude differences.

### G. Representational Similarity Analysis

While node-strength analysis focuses on local topological hubs, understanding the overall representational geometry requires a broader metric. Therefore, Representational Similarity Analysis (RSA) was performed to evaluate whether the global internal representation of the model captures the network dynamics reflected in the effective connectivity patterns [46].

The results in Table VI and Fig. 5B show that the RSA scores for the proposed preprocessing pipeline are consistently higher than the baseline across almost all connectivity methods and frequency bands. This positive difference indicates a tendency for the proposed pipeline to yield representations that are structurally closer to the underlying EEG connectivity patterns.

However, it is crucial to note that none of these increases reached statistical significance after FDR correction (all *p >* 0.05). Despite this lack of strict statistical significance, several conditions, particularly within the GPDC method across broadband and higher frequencies, exhibited moderate effect sizes (Cohen’s *d* reaching up to 0.66).

In terms of brain activity, this suggests that the proposed preprocessing introduces a subtle positive shift in the model’s ability to capture global representation structures associated with visual area communication dynamics, even though the absolute magnitude of this difference remains relatively small [43]. Ultimately, this confirms that the global network representations in both pipelines successfully preserve and reflect similar fundamental brain network activity patterns during visual stimulus processing in the RSVP paradigm [20], [38].

## V. Conclusion

This study systematically evaluated how preprocessing strategies shape channel-wise attention representations and their biological interpretability in Transformer-based visual EEG decoding. Our findings demonstrate that rigorous EEG preprocessing, specifically the integration of ICA and notch filtering, successfully suppresses non-neural artifacts, such as frontal ocular contamination and line noise, without significantly compromising decoding accuracy or model resilience to external signal interference. The stable cross-generalization observed across preprocessing conditions suggests that the removed spectral components are not essential for task performance. However, this result does not constitute direct evidence that the model relies solely on neural activity, as the learned representations may adapt to preprocessing-induced changes in the signal structure.

Crucially, our spectral and temporal ablation analyses revealed that the Transformer’s decoding capability is predominantly driven by low-frequency oscillations (specifically the theta and delta bands) and early post-stimulus intervals (0 to 400 ms). This spatiotemporal dependency strongly mirrors the known neurophysiological markers of early perceptual and attentional processing within rapid visual presentation paradigms.

Furthermore, biological interpretability appears to be metric-dependent in this study: preprocessing left hub-level alignment broadly stable while producing a modest positive shift in RSA-based global representational correspondence. While Representational Similarity Analysis (RSA) indicated that the proposed pipeline introduced subtle global structural alignments with directed connectivity metrics like GPDC, the core topological organization of the network (evaluated via node-strength correlation) remained highly stable across both pipelines. Rather than universally increasing interpretability in a linear fashion, extensive preprocessing refines the specific structural information captured by the attention mechanism without altering its overall biological plausibility. Ultimately, this work provides a balanced framework for bridging data-driven machine learning optimization with neurophysiological consistency, laying the groundwork for developing more transparent and medically trustworthy brain-computer interfaces.

